# Programmable single and multiplex base-editing in *Bombyx mori* using RNA-guided cytidine deaminases

**DOI:** 10.1101/269969

**Authors:** Yufeng Li, Sanyuan Ma, Le Sun, Tong Zhang, Jiasong Chang, Wei Lu, Xiaoxu Chen, Yue Liu, Xiaogang Wang, Run Shi, Ping Zhao, Qingyou Xia

**Author notes:** These two authors contributed equally to this work (Yufeng Li, Sanyuan Ma). Correspondence: Qingyou Xia.

## Abstract

Standard genome editing tools (ZFN, TALEN and CRISPR/Cas9) edited genome depending on DNA double strand breaks (DSBs). A series of new CRISPR tools that convert cytidine to thymine (C to T) without the requirement for DNA double-strand breaks were developed recently, which have changed this status and have been quickly applied in a variety of organisms. Here, we demonstrate that CRISPR/Cas9-dependent base editor (BE3) converts C to T with a high frequency in the invertebrate *Bombyx mori* silkworm. Using BE3 as a knock-out tool, we inactivated exogenous and endogenous genes through base-editing-induced nonsense mutations with an efficiency of up to 66.2%. Furthermore, genome-scale analysis showed that 96.5% of *B. mori* genes have one or more targetable sites being knocked out by BE3 with a median of 11 sites per gene. The editing window of BE3 reached up to 13 bases (from C1 to C13 in the range of gRNA) in *B. mori*. Notably, up to 14 bases were substituted simultaneously in a single DNA molecule, with a low indel frequency of 0.6%, when 32 gRNAs were co-transfected. Collectively, our data show for the first time that RNA-guided cytidine deaminases are capable of programmable single and multiplex base-editing in an invertebrate model.

## INTRODUCTION

The clustered regularly interspaced short palindromic repeat (CRISPR/Cas9) has been widely used for site-specific genome editing in various organisms and cell lines(SANDER and JOUNG 2014). Under the direction of a guide RNA (gRNA), the Cas9 nuclease binds to an opened DNA strand that paired with the gRNA and induces a double-strand break (DSB). At this point, the intrinsic cellular DNA-repair mechanism will generally repair the DSB via non-homologous end joining (NHEJ), resulting in deletions or insertions (indels) in most cases. When present, the cell can use a homologous DNA fragment as a template to repair the DSB by another mechanism, known as homology-directed repair (HDR). The efficiency of NHEJ-induced indel formation is always higher than that of HDR-mediated gene correction, although various efforts have been made to increase the frequency of HDR(CHU *et al.* 2015; MARUYAMA *et al.* 2015; PAQUET *et al.* 2016; RICHARDSON *et al.* 2016; SAKUMA *et al.* 2016). ZFN, TALEN, and CRISPR/Cas9 all rely on the generation of a DSB, which can lead to many potential defects such as unexpected indels, off-target cleavage, and decreased cell proliferation when targeted to copy number-amplified (CNA) genomic regions (SHEN and IDEKER 2017). Thus, approaches to precisely edit genome avoiding DSBs are needed.

Several modified systems have been developed to overcome the drawbacks of DSB-based genome editing. Komor et al. showed that the rat cytidine deaminase rAPOBEC1 could be linked to nCas9 (Cas9 nickase) to efficiently substitute C with T at target sites without generating DSBs (KOMOR *et al.* 2016). After three generations of modification, they developed the final version of base editor (BE), named BE3, which showed up to a 74.9% mutation efficiency in mammalian cells, that was composed of rAPOBEC1, nCas9 (A840H), and UGI (uracil DNA glycosylase inhibitor). Kondo et al. engineered dCas9/nCas9 and the activation-induced cytidine deaminase (AID) ortholog PmCDA1 to assemble a complex (Target-AID) that performed highly efficient specific point mutations (NISHIDA *et al.* 2016). In a parallel study, Ma et al. fused UGI with dcas9-AIDx (AICDA-P182X) to convert targeted cytidine specifically to thymine, in a process referred to as targeted AID-mediated mutagenesis (MA *et al.* 2016). Additionally, to further optimize the base editor BE3, a series of studies were conducted to expand the number of target sites, narrow the width of the editing window (KIM *et al.* 2017c), improve the efficiency and product purity (KOMOR *et al.* 2017), and reduce off-target effects (KIM *et al.* 2017a). For base editor BE3 converts C:G base pairs to T:A base pairs with a high efficiency, several groups have used BE3 to silence genes through base-editing-induced nonsense mutations (BILLON *et al.* 2017; KIM *et al.* 2017b; KUSCU *et al.* 2017).

The development of base-editing systems has both improved the scope and effectiveness of genome editing. Site-directed mutagenesis in the genome can be achieved by design, rather than randomly achieved through wild-type Cas9. To date, several organisms have been subjected to base editing, mostly using BE, including mammalian cells (KOMOR *et al.* 2016; MA *et al.* 2016), rice (LU and ZHU 2017; ZONG *et al.* 2017), wheat (ZONG *et al.* 2017), tomatoes (SHIMATANI *et al.* 2017), mice (KIM *et al.* 2017b), and zebrafish (ZHANG *et al.* 2017). Komor et al. corrected two disease-relevant mutations that cause Alzheimer’s disease in mammalian cells by BE3. Zhang et al. converted a proline residue at position 302 in the *tyr* gene to serine, threonine, or alanine to mimic the oculocutaneous albinism (OCA) mutation in zebrafish by BE3. However, no reports have described the performance of any base-editing system in invertebrates.

*Bombyx mori* (silkworm) is a model organism for studying invertebrate biology and an economically important lepidopteran insect (GOLDSMITH *et al.* 2005). The genome-editing tools ZFN, TALEN, and CRISPR/Cas9 have all been applied in *B. mori* (MA *et al.* 2012; LIU *et al.* 2014; MA *et al.* 2014a; MA *et al.* 2014b; MA *et al.* 2014c; LIU *et al.* 2017). Here, we demonstrated that BE3 performed base editing by targeting the *Blos2* and *Yellow-e* genes with efficiencies of 25% and 51.2% in *B. mori*, respectively. Then, we demonstrated that BE3 could efficiently knock out exogenous and endogenous genes through base-editing-induced nonsense mutations with efficiencies of up to 66.2%. Furthermore, we found 151,551 targetable knockout sites in genome-wide. The deamination window of BE3 in *B. mori* spans from C1 to C13 within the protospacers. In addition, by co-transfecting 32 gRNAs and BE3 simultaneously, we generated up to 14 C:G base-pair substitutions in one DNA molecule, with few indels observed.

## RESULTS

### Establishing a base-editing system for *B. mori*

To confirm whether base editing could be achieved by BE3 (rAPOBEC1-XTEN-nCas9-UGI) in *B. mori*, we first constructed the BE3 vector. DNA encoding rAPOBEC1, the XTEN linker, a partial nCas9 sequence with an A840H mutation, and UGI was commercially synthesized, using *B. mori* codon-optimized sequences, and then was inserted into the pUC57-T-simple plasmid. The Hsp70 promoter and SV40 terminator sequence (MA *et al.* 2017) were used to initiate and terminate BE3 expression, respectively. The complete BE3 vector was constructed by inserting the fragment encoding both Cas9 and nCas9 (A840H) into the synthesized plasmid after the digestion with *Nde*I and *Bam*HI (**Figure 1A**). Then, the BE3 vector was transfected into *B. mori* embryo cell line, BmE. Cellular proteins were extracted at 3 days post-transfection. Expression of the BE3 protein was confirmed by western blot analysis (**Figure 1B**). To evaluate the efficacy of BE3-based genome editing in invertebrate cells, two previously constructed gRNAs targeting *Blos2* and *Yellow-e* in *B. mori* cells were selected (LIU *et al.* 2014). Based on the typical editing window *in vitro* (KOMOR *et al.* 2016), ranging from position 4 to 8 in the protospacer counting from the distal end to the protospacer-adjacent motif (PAM), both gRNAs have C located in position 5 (C5) and 7 (C7). We co-transfected gRNAs and BE3 vectors into the BmE cells, and cellular genomic DNA was extracted after 3 days without selection. Genomic regions spanning the target sites were amplified by PCR for sequencing. Sanger sequencing chromatograms for both the *Blos2* and *Yellow-e* PCR products showed a set of C/T peaks in the target sites, indicating that the base substitutions did occur (**Figure 1C, D**). To confirm the base substitutions, the PCR products were cloned into the pEASY-T5 vector by T-A cloning. Sanger sequencing of individual clones showed different types of mutations in the DNA sequences. For *Blos2*, two out of five clones (40%) had C-T substitutions. The efficiency of C-T substitutions for *Yellow-e* reached 51.2% in 41 examined clones. Among the studied positions (C5, C7, C9, and C11), C5 showed the highest mutation rate. The observation of base substitutions in C9 and C11 was beyond our expectation because C9 and C11 laid outside of the typical editing window observed *in vitro* and in plants (KOMOR *et al.* 2016; ZONG *et al.* 2017). Taken together, these results demonstrated that the BE3–gRNA nuclease complex effectively and site-specifically substituted C with T in invertebrate *B. mori* cells. Notably, the base-editing window of BE3 in *B. mori* may be wider than that *in vitro* and plants.

**Figure 1.**
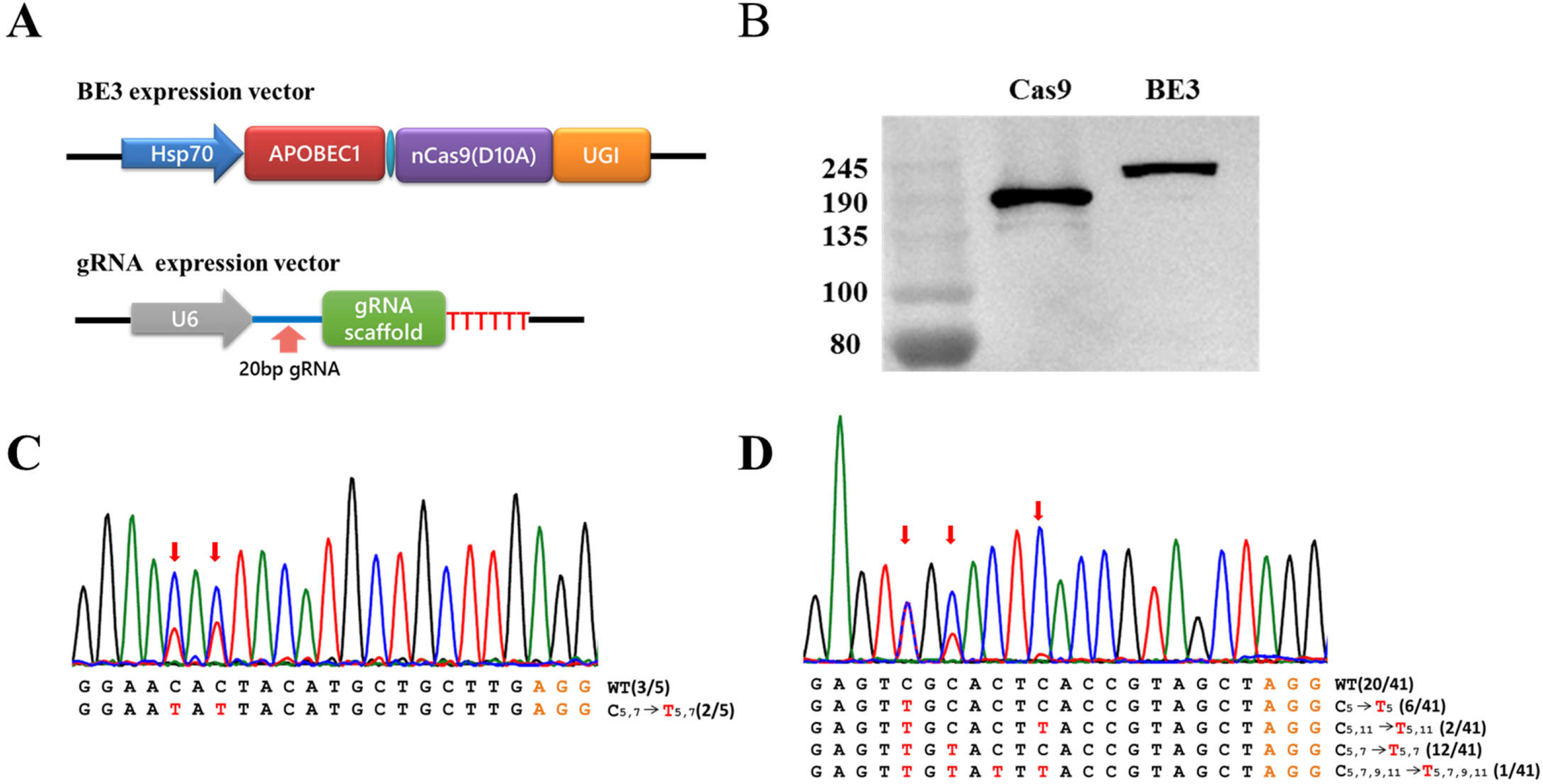
Base editing in *B. mori.* (**A)** Schematic representation of the BE3- and gRNA-expression vectors. (**B)** Detection of BE3 expression in the BmE cell line by western blotting. (**C, D**) Sanger sequencing of targeted genomic regions of *Blos2* and *Yellow-e*. *Red arrows* point to overlapping peaks. The orange and red letters mark PAM sites and converted bases, respectively. The numbers in trumpet font indicate the position of C or T.

### Using BE3 to knock out the exogenous genes *mCherry* and *Puromycin* in *B. mori* cells

To expand the application of BE3 in *B. mori*, we evaluated BE3 as a knock-out tool through inducing premature stop codons (TAG, TGA, or TAA) by converting C:G base pairs to T:A base pairs for four codons (CAA, CAG, CGA, TGG) in coding strands. Four gRNAs with the potential of generating stop codons in the exogenous *mCherry* reporter gene were designed (**Table S1**). We co-transfected these four gRNAs (mCherry-1, 2, 3, or 4) together with BE3 individually into a transgenic cell line (BmE-mCherry), which harbors the *mCherry* expression cassette (unpublished). At 12 days post-transfection, we evaluated the knock-out efficiency by measuring the average fluorescence intensity of mCherry via flow cytometry. Significantly decreased mCherry signal was found with all four gRNAs compared with control (**Figure 2A**, **Figure S1**). We also observed markedly decreased mCherry-positive cells in cells harboring the BE3‒gRNA complex by fluorescence microscopy (**Figure 2A**), in accord with the flow cytometry results. However, it should be noted that the most effective gRNAs were mCherry-2 and mCherry-3 (59.1 and 66.2%, respectively), rather than mCherry-1, which targeted the 5′-most mCherry sequence among the four gRNAs. These results may have been observed because mCherry-2/3 targeted TGG with two potentialities of inducing stop codons. To further confirm that mCherry was knocked out through a substitution-induced nonsense mutation, we amplified and sequenced the target genomic regions. Sanger sequencing of both PCR products and individual clones showed several base substitutions, resulting in stop codon mutations (**Figure 2B**). Although no mutations were found with mCherry-2 among 10 clones, the Sanger sequencing chromatogram for the PCR product did show a overlapping peak at the target site, where CAG was converted to the stop codon TAG (**Figure 2B**), resulting in *mCherry* being knocked out. To test the universality of BE3 in *B. mori*, we then designed two gRNAs (Puromycin-1, 2) targeting another exogenous gene *Puromycin*. Cellular proteins were extracted at 12 days post-transfection. Western blot results indicated that *Puromycin* protein levels dramatically decreased after transfection with either gRNA (**Figure 3A**). Sanger sequencing of PCR products and clones also showed the conversions of C:G to T:A. The stop codons TAA and TGA generated by BE3 caused *Puromycin* to be knocked out with efficiencies of 58.3% (7/12) and 27.2% (3/11), respectively (**Figure 3B**). Taken together, our data indicated that we effectively knocked out the exogenous genes *mCherry* and *Puromycin* via substitution-induced nonsense mutations mediated by BE3 in *B. mori*.

**Figure 2.**
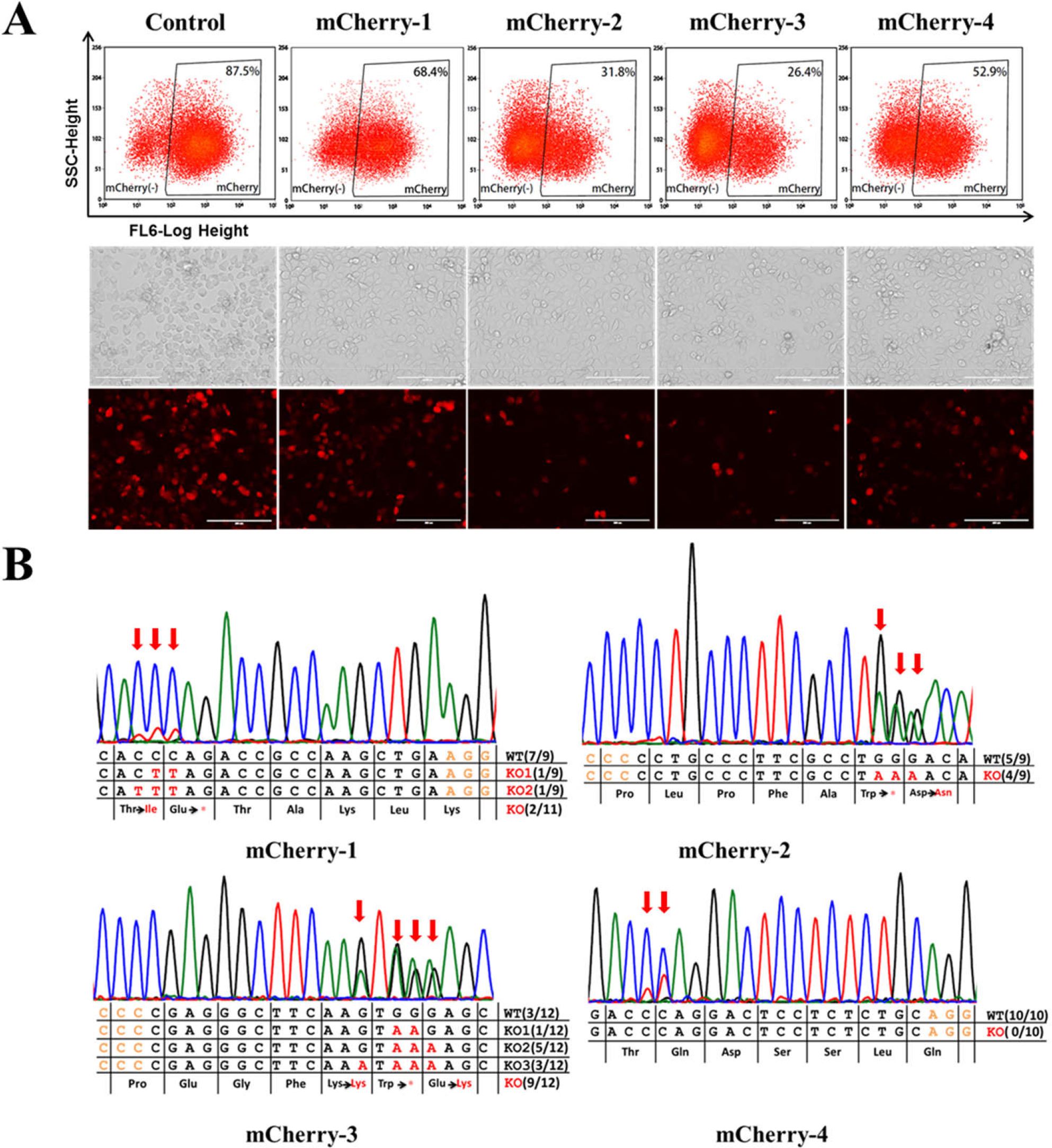
Using BE3 to knock out the *mCherry* reporter gene by introducing a premature stop codon. (**A**) Flow cytometry-based quantification of mCherry expression at 12 days post-transfection (upper panel). Light and fluorescence images show decreased mCherry-positive cells (lower panel). (B) Sanger sequencing results show the base conversions in targeted genomic regions of four *mCherry* gRNAs. The encoded amino acids are shown below. Asterisks in red represent stop codons.

**Figure 3.**
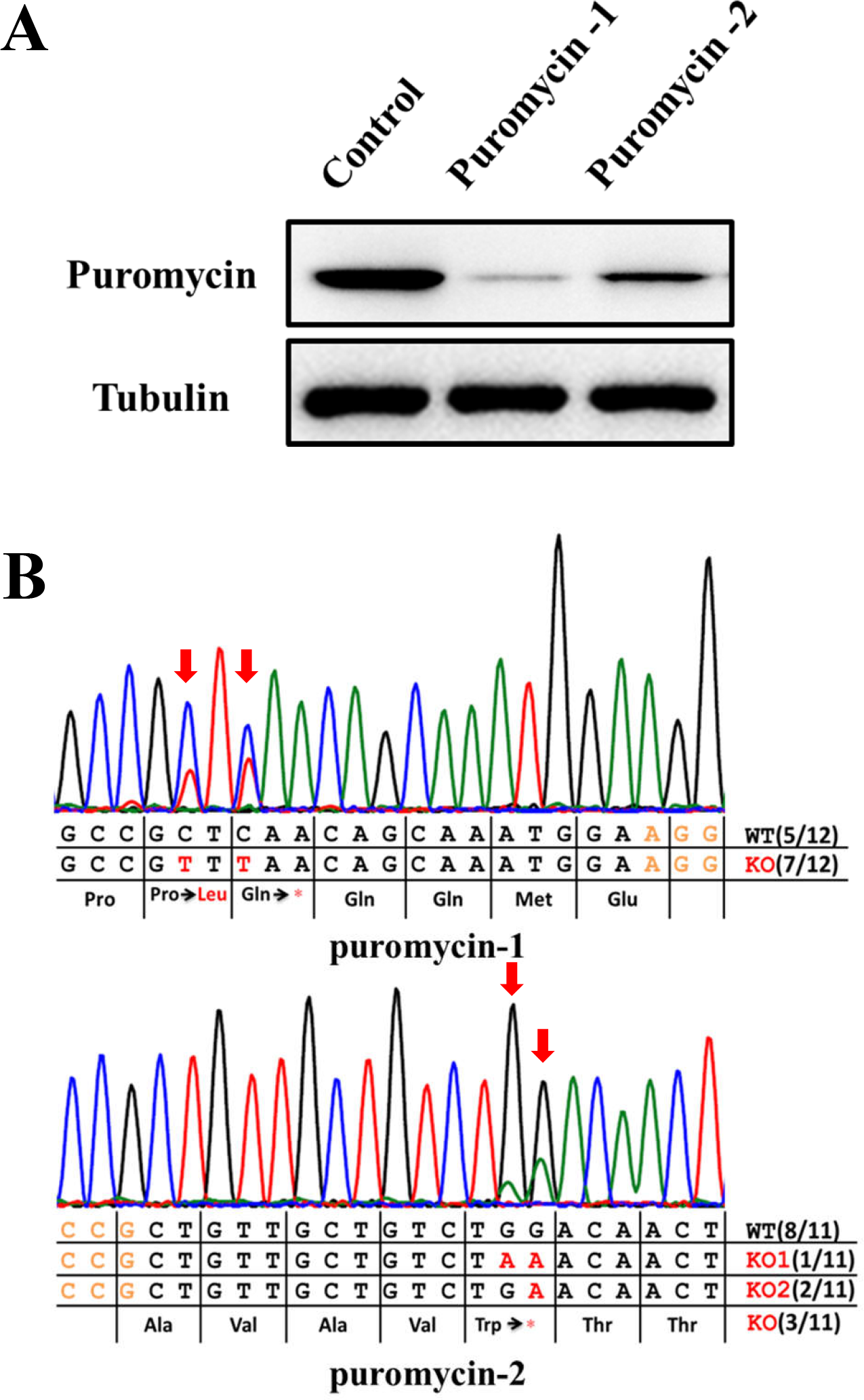
Using BE3 to knock out the exogenous *Puromycin* gene by introducing a premature stop codon. (**A**) Sanger sequencing show the base conversions in targeted genomic regions of two *Puromycin* gRNAs. (**B**) Protein levels were analyzed by western blotting at 12 days post-transfection. Alpha-tubulin was detected as a control protein.

### Using BE3 to knock out the endogenous genes *BmGAPDH* and *BmV-ATPase B*, and predicting knocked out loci on a genome-wide scale

To determine whether BE3 could generate effective substitution-induced nonsense mutations for endogenous genes, four gRNAs were designed to target *BmGAPDH* (GAPDH-1, 2) and *BmV-ATPase B* (ATPase-1, 2). We extracted cellular proteins at 12 days post-transfection. Western blot analyses indicated that protein levels noticeably decreased for GAPDH-1 and ATPase-1, compared with control (**Figure 4A**). Sanger sequencing chromatograms for four PCR products showed overlapping peaks in the target sites, that where nonsense mutations arose (**Figure 4B**), causing *BmGAPDH* and *BmV-ATPase B* to be knocked out. The Sanger sequencing results were consistent with the western blot results. Collectively, these findings showed that BE3 could mediate effective substitution-induced nonsense mutations for endogenous genes in *B. mori* cells. However, not all gRNAs work exceedingly well with BE3 in knocking out genes (as also found with Cas9), such as GAPDH-2 and ATPase-2 (**Figure 4A, B**).

**Figure 4.**
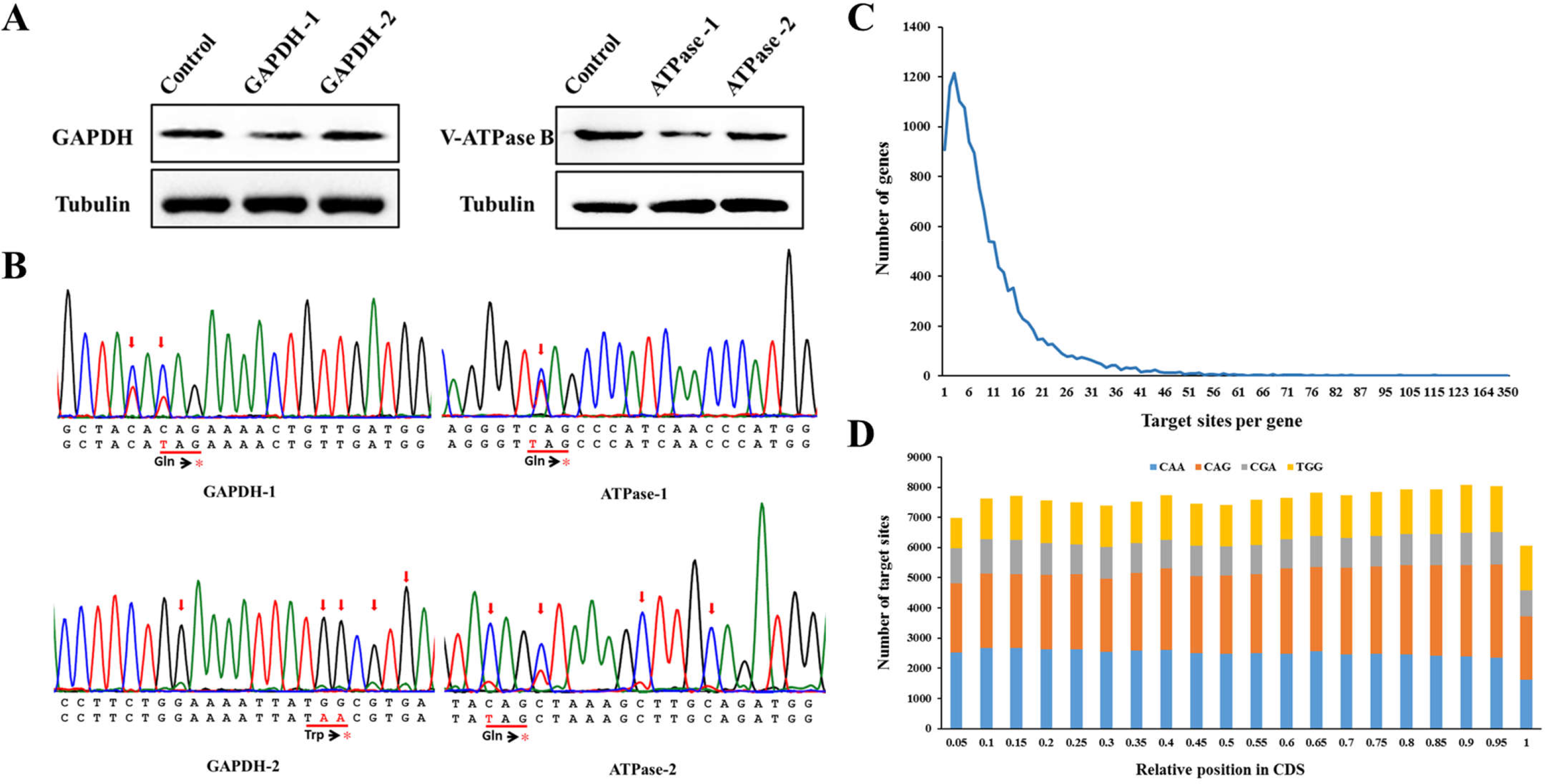
Using BE3 to knock out the endogenous *GAPDH* and *V-ATPase B* genes by introducing a premature stop codon. (**A**) Protein levels were analyzed by western blotting at 12 days post-transfection. Alpha-tubulin was detected as a control protein. (**B**) Sanger sequencing of genomic regions targeted by *GAPDH* and *V-ATPase B* gRNAs. The encoded amino acids are shown below. Asterisks in red represent stop codons. (**C**) The number of genes with different numbers of targetable knockout sites per gene. (**D**) Relative positions of knockout sites in CDSs. Four targetable codons are shown in different colors.

To determine the targetable sites for knock-out by BE3 at the genome scale, we identified all candidate codons (CAA, CAG, CGA and TGG) that can potentially be converted to stop codons (TAA, TAG, TGA) by BE3. This genome-scale analysis revealed a pool of 151,551 targetable knockout sites in 14,106 CDSs, with a median of 11 sites per gene and 96.5% targetable genes (among 14,623 total genes) (**Figure 4C**). Furthermore, the distributions of these codons were well-distributed within the CDSs, suggesting that approximately half of these targetable sites could stop mRNA translations within the first 50% of their encoded protein sequences (**Figure 4D**).

### The editing window of BE3 is mostly in C4–C7, although effective C substitutions can occur within C1–C13

Sanger sequencing for previous T clones showed that base substitutions could be generated at C3‒C9 and C11. However, these results may not fully represent the editing window of BE3 in *B. mori.* Next, we designed 32 gRNAs to target the enhanced green fluorescent protein (EGFP) gene within the 5′ 400 base pairs (bp) (**Figure 5A**). Each gRNA was co-transfected with the BE3 vector into the BmE-EGFP cell line (a transgenic cell line harboring an EGFP-expression cassette, unpublished). The PCR products for targeted genomic regions were sequenced by next-generation sequencing (NGS). We averagely obtained 96,782 DNA reads for every gRNA. The efficiencies of 32 gRNAs, ranging from 3.4% to 25.3%, were calculated by counting every DNA read with one or more C:G to T:A substitutions within the 20-bp gRNA target sites (**Figure 5B**). The indel frequencies of the 32 gRNAs were ≤5.0%, except for gRNA21 (8.1%), while the control showed a 1.2% indel frequency (**Table S3**). Then, we analyzed the efficiency of each base substitution for each gRNA in detail (**Figure 5C**). The different positions of C:G base pairs in the range of 32 gRNAs probably reflected diverse efficiencies (**Table S1**). For each C:G base pair located at different positions within each gRNA and for each gRNA had a different number of C:G base pairs, we counted the number of gRNAs with valid Cn:Gn substitutions and the number of gRNAs that have corresponding Cn:Gn. The ratio of these two numbers indicated the probability of each C:G substitution within 20-bp gRNA target site (**Figure 5D**). Thus, now we have a clear understanding of the editing window which is mostly within C4-C7 although effective C substitutions spread C1-C13 (**Figure 5C**).

**Figure 5.**
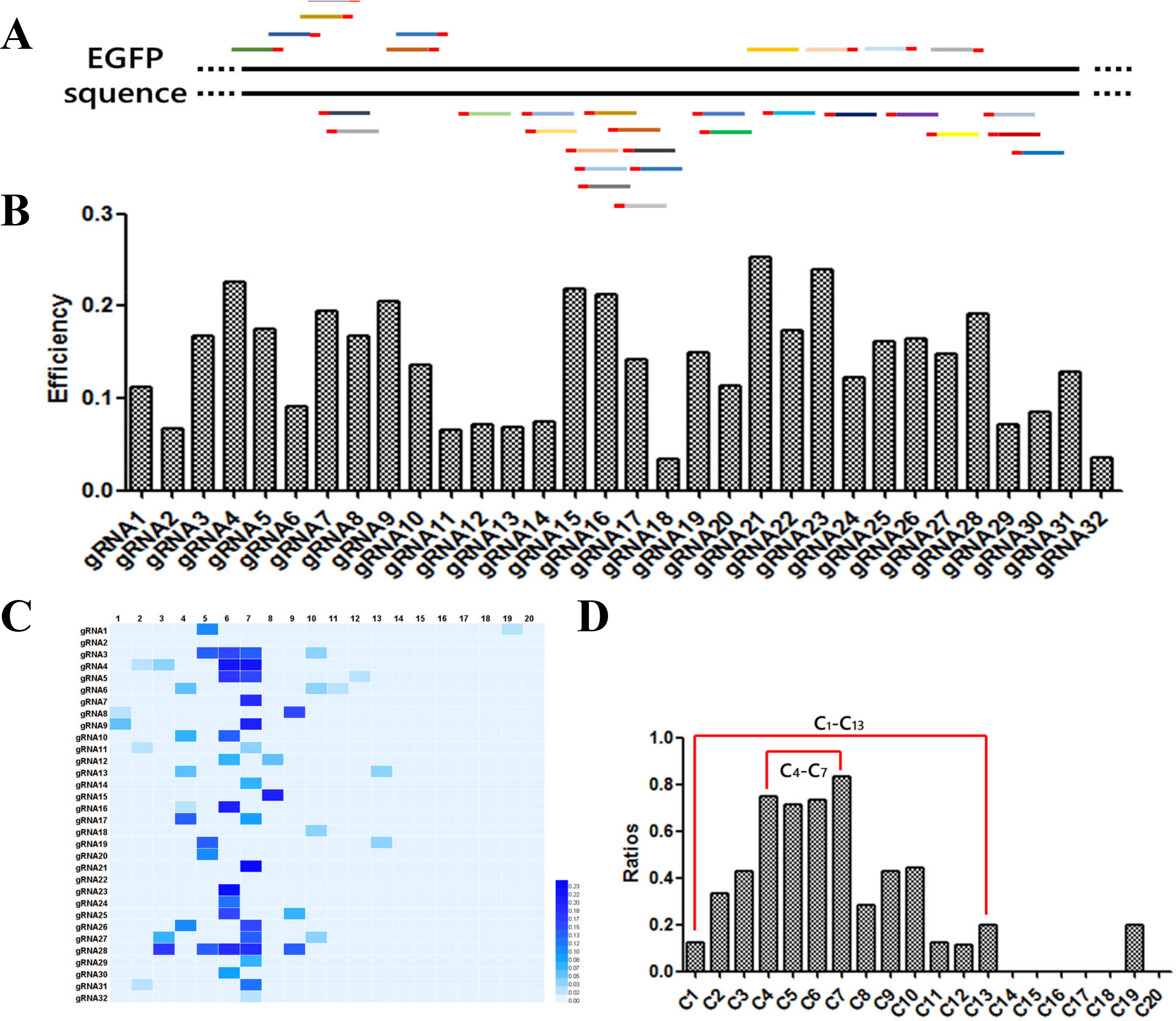
The efficiencies of 32 EGFP gRNAs and the editing window of BE3 in *B. mori*. **(A**) Schematic representation of 32 gRNAs targeting EGFP. (**B**) Bar graphs show the efficiencies of the 32 gRNAs designed to target EGFP. (**C**) The heatmap shows the efficiency of each base substitution in each gRNA. The darker of the colors are, the more efficient of the base substitutions are. (**D**) Bar graphs show that the editing window of BE3 in *B. mori* ranges from C1 to C13.

### Multiple base-editing with an extremely low indel frequency by BE3

Previous studies have indicated that DSBs generated by Cas9 might lead to excessive DNA damage that could cause cell death (SHEN and IDEKER 2017) and targeted chromosome elimination (ZUO *et al.* 2017) when targeting several genomic loci simultaneously. However, BE3, which is much more temperate than Cas9, with respect to the genome, demonstrated base editing without generating DSBs. Thus, BE3 should be more suitable for performing multiple genome edits, without causing excessive DNA damage and indels (KOMOR *et al.* 2016; KIM *et al.* 2017a; ZONG *et al.* 2017). All 32 gRNAs that target EGFP were co-transfected together with the BE3 vector into cells. The PCR products for the targeted genomic regions were sequenced by NGS. We found a range of mutation types (1–14 base substitutions) with an extremely low indel frequency of approximately 0.6%, which was the equal to the control (**Figure 6A**, **Table S3**). The more base substitutions generated by BE3 simultaneously, the fewer reads that were observed (**Figure 6A**). We further investigated these types of mutation sequences, and found that several C:G base pairs that located in different positions were mutated to T:A base pairs simultaneously in a single DNA read (**Figure 6B**). Although only a few reads showed 10 or more base-substitutions in one DNA read, they did not exist in the control. Together, our data indicate that BE3 represents an improvement over the Cas9 nuclease in inducing multiple base mutations with an extremely low indel frequency and no DSBs.

**Figure 6.**
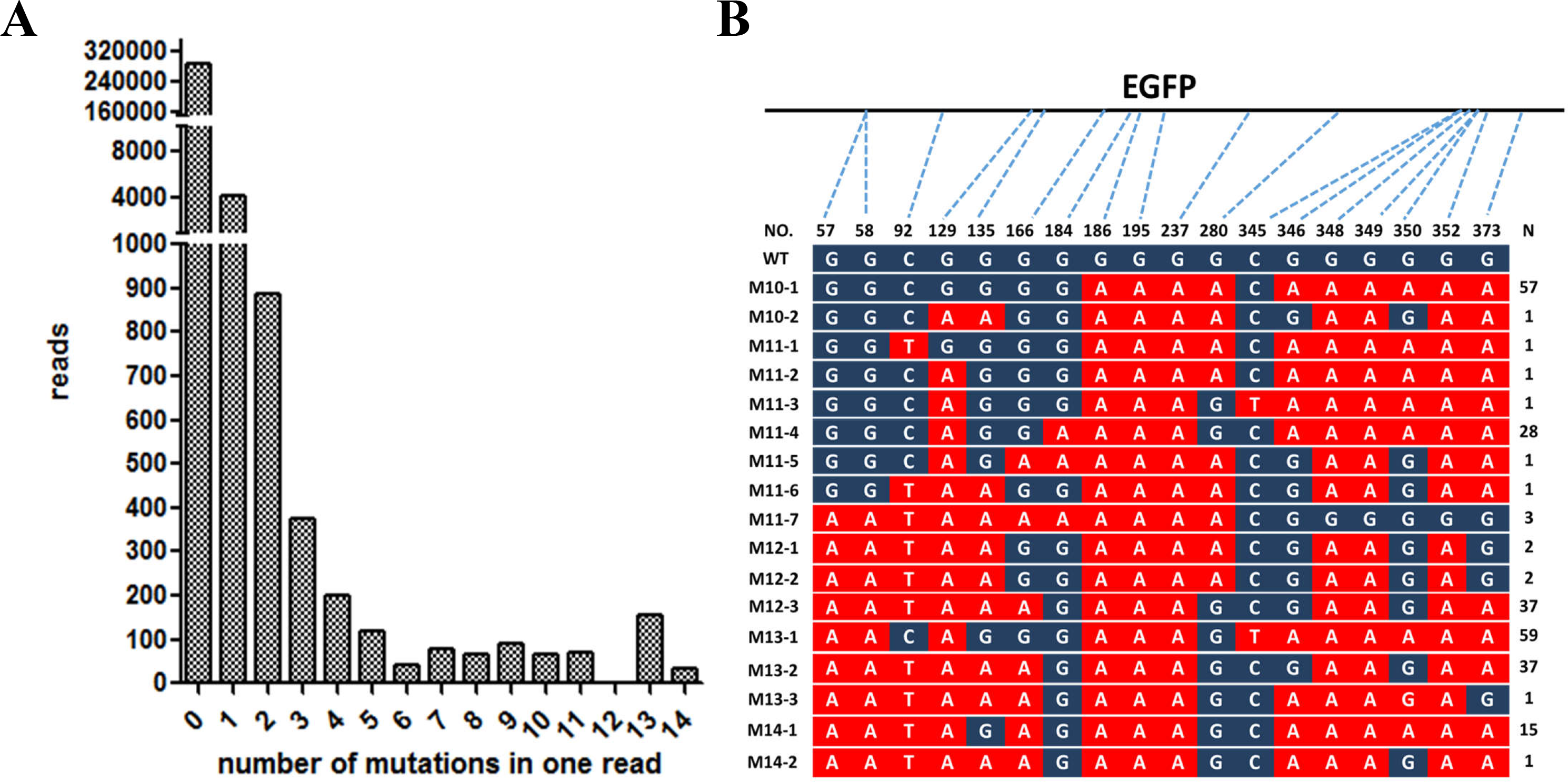
Multiple base substitutions generated by BE3. **(A)** Bar graphs show the reads of multiple mutations after co-transfected 32 gRNA vectors and the BE3 vector. (**B**) Schematic representation of 10 or more base substitutions in a single DNA read. The converted bases are marked in red. n1: number of base substitutions. n2: reads of DNA with multiple base substitutions

## DISCUSSION

In recent years, the CRISPR/Cas9 genome-editing technology has been rapidly and widely adapted in cutting-edge fields such as stem cell biology, genomic biology, developmental biology, and cancer research because of its high efficiency and simplicity (HSU *et al.* 2014). However, the existing CRISPR/Cas9 system does not meet the needs of gene therapy sufficiently for human genetic diseases caused by point mutations. Advances and developments in base-editing tools show great potential for maintaining the power of genome editing, while reducing its uncertainty and risk. By fusing nCas9 with cytidine deaminases, scientists have used BE3 to generate site-specific base conversions from C:G base pairs to T:A base pairs without inducing DSBs and effectively avoid indels. In addition, a series of subsequent reports have described improvements in its performance and generating nonsense mutations (BILLON *et al.* 2017; KIM *et al.* 2017b; KUSCU *et al.* 2017).

Recently, BE3 was confirmed to function stably and efficiently in a variety of organisms. The efficiencies were up to 74.9%, 43.48%, and 28% in mammalian cells (KOMOR *et al.* 2016), plants (ZONG *et al.* 2017) and zebrafish (ZHANG *et al.* 2017), respectively. Here we report that BE3 can mediate the conversion of cytidine to thymine with an efficiency of up to 66.2% in *B. mori*, which to our knowledge, is the first report demonstrating BE3-mediated base editing in an invertebrate species. Previous reports have shown the successful use of BE3 as a knock out tool with high efficiency in mouse embryos and mammalian cells (BILLON *et al.* 2017; KIM *et al.* 2017b; KUSCU *et al.* 2017). Here, we also used BE3 to efficiently inactivate exogenous and endogenous genes through substitution-induced nonsense mutations in *B. mori*. A recent report showed that the plant base editor (nCas9-PBE) could induce C to T with deamination windows covering seven bases from C3–C9 in the protospacers (ZONG *et al.* 2017), whereas we detected a broader editing window with effective C substitutions spreading from C1 to C13.

Genome editing without generating DSBs is the biggest advantage of the base-editing system over Cas9. Such a capacity will guarantee the integrity of the genome to a much larger extent. In this study, we only detected a few indels generated by 32 gRNAs individually targeting EGFP with an average indel frequency of 2.2%, compared with a 1.2% indel frequency in control samples (which might have been induced by PCR or sequencing) (**Table S3**). Previous data showed that the indel frequency for BE3 was also very low in mammalian cells and plants (KOMOR *et al.* 2016; ZONG *et al.* 2017), but that is different in *B. mori*. We suspect that the frequency of indels may be diverse because the efficiency of BE3 and mechanisms of DNA-damage repair differ between various organisms. However, when we co-transfected all 32 gRNAs together with the BE3 vector, the indel frequency in experimental group was almost the same as that in control cells. This result indicated that multiple bases can be edited simultaneously with almost no indels occurring around the target sites.

Using the CRISPR/Cas9 system and a pool of gRNAs, two groups (KATHERINE 2016; GARST *et al.* 2017) mapped the functional domains of proteins and promoters through error-prone NHEJ or correction-prone HDR. However, the unavoidable defects of both approaches were that indels were caused by NHEJ or HDR, resulting in genome damage. Due to its DSB-free nature, BE3 is more useful for identifying alterations conferring a gain of function to modified proteins. Our findings showed that, with 32 gRNAs together, 14 C:G to T:A conversions occurred in EGFP with a indel frequency that is comparable to control. Using the C:G base editor BE3 and the recently described A:T base editor ABE (GAUDELLI *et al.* 2017), all four transition mutations can be programed with a library of gRNAs to enable mapping of thousands of parallel amino acids and promoter mutations simultaneously. Additionally, the limits associated with the PAM sites can be significantly relieved by the development of various Cas9 enzymes that recognize other than NGG (KLEINSTIVER *et al.* 2015a; KLEINSTIVER *et al.* 2015b; RAN *et al.* 2015; GAO *et al.* 2017).

In summary, we demonstrated site-specific base editing by BE3 with high efficiency in an invertebrate organism, for the first time. This study may help in generating efficient base-editing systems in other organisms. The editing window, ranging from C1 to C13, is wider than that in plants (ZONG *et al.* 2017), which provides greater potential and feasibility for modifying the genome at target sites with a larger scope. We also used BE3 to knock out exogenous and endogenous genes with high efficiency through substitution-induced nonsense mutations in *B. mori*. The frequency of introducing a stop codon was up to 66.2%, which was much more efficient than that found with SpCas9, SaCas9, and Cpf1 in *B. mori* cells (LIU *et al.* 2014; MA *et al.* 2017). From this point of view, BE3 is more suitable for knock-out studies in invertebrates. Moreover, genome-wide bioinformatics analysis revealed 151,551 targetable knockout sites located in 96.5% of all *B. mori* genes, with approximately 11 sites per gene. BE3 provides an alternative for functional-genomics studies and other knock-out experiments and represents an improvement over Cas9 in terms of its ability to perform multiple base substitutions, without generating DSBs. Using 32 gRNAs simultaneously, we found up to 14 base mutations occurred in EGFP, with scarcely any indels. This result illustrates a novel strategy that using a pool of gRNAs and BE3 offers an enormous potential for mapping functional protein domains by generating diverse variants via BE3-mediated mutagenesis. Hence, BE3 is more applicable for identifying gain-of-function mutations in proteins. This is also a good news because specific modifications via Cas9-mediated HDR have been difficult to achieve in many organisms. This new CRISPR tool for base editing should have a substantial impact on basic researches and the gene therapy for clinical treatments.

## MATERIALS AND METHODS

### Design and construction of the BE3- and gRNA-expression vectors

DNA encoding rAPOBEC1, the XTEN linker, partial nCas9 with the A840H mutation, and UGI was *B. mori* codon-optimized and synthesized by GenScript service, and then inserted into the pUC57-T-simple plasmid. The complete BE3 vector was constructed by inserting the same fragment of Cas9 and nCas(A840H) into the synthesized plasmid after digestion of *Nde*I and *Bam*HI (New England BioLabs). The gRNA-expression vector pUC57-gRNA was described previously (LIU *et al.* 2014). gRNA sequences (Table S1) were synthesized as two complementary oligonucleotides, annealed, and inserted into the pUC57-gRNA plasmid after *Bbs*I digestion.

### Cell culture and transfection

The *B. mori* embryo cell lines BmE, BmE-mCherry, and BmE-EGFP was established in our laboratory and maintained at 27°C in Grace insect medium (Gibco, Thermo Fisher) containing 10% fetal bovine serum (Gibco, Thermo Fisher). Cells were seeded in 12-well plates (Corning) and transfected at approximately 80% confluency. Plasmids (1.8 μg total) were transfected at a 1: 1 ratio into cells using the X-tremeGENE HP DNA Transfection Reagent (Roche), following the manufacturer’s recommended protocol.

### Flow cytometry and fluorescence imaging

Cells were harvested at 12 days post-transfection. A MoFlo XDP flow cytometer (Beckman) was used to measure mCherry fluorescence, and Summit software was used to analyze the data. In addition, light and fluorescence microscopy images were captured using an EVOS FL Auto microscope (Life Technologies).

### Protein extraction and western blot analysis

Cellular proteins were extracted using NP-40 lysis buffer (Beyotime) on day 12 post-transfection. Proteins were quantified using a BCA Kit (Beyotime). Equal amounts of proteins were resolved by 12% sodium dodecyl sulfate-polyacrylamide gel electrophoresis and transferred to a polyvinylidene fluoride (PVDF) membrane. Next, 5% milk was used to block the PVDF membranes, which were then incubated with primary antibodies (diluted 1: 1,000 in 1% milk) for 2 h. Subsequently, the PVDF membranes were incubated with anti-mouse or anti-rabbit secondary antibodies (diluted at 1:10,000 in 1% milk) for 1 h. Membranes were developed with Thermo Fisher ECL reagent and then imaged with a western blot processor.

### Genome-wide analysis of knock-out sites

To identify knock-out sites on a genome-wide scale, we first retrieved all CDSs for *B. mori* from the Silkworm Genome Database. gRNAs and the nucleotide motifs CAG, CGA, CAA, and TGG were searched using the fuzznuc EMBOSS explorer. Based on the editing window of BE3 for *B. mori*, nucleotide motifs (with a multiple of three for the last base) at position 1‒13 within each gRNA were selected using the intersectBed and shell script. These nucleotide motifs were defined as potential knock-out sites.

### Sanger sequencing

Genomic DNA extracted using the E.Z.N.A. Tissue DNA Kit (Omega) following the manufacturer’s recommended protocols after cells were transfected for 3 days. The genomic regions of interest were amplified using site-specific primers (Table S2). PCR products were generated with PrimerSTAR Max DNA Polymerase (Takara) according to the manufacturer’s instructions, then loaded on a 2% agarose gel. The Gel Extraction Kit (Omega) and pEASY-T5 vector (Transgene) were used for PCR product purification and T-A cloning, respectively. The PCR and T clone products were then subjected to Sanger sequencing.

### NGS experiments

Thirty-two gRNA vectors targeting EGFP were co-transfected individually with the BE3 vector. Cellular genomic DNA was extracted using the E.Z.N.A. Tissue DNA Kit (Omega) following the manufacturer’s protocols at 3 days post-transfection. The target regions were amplified using site-specific primers with added barcodes. Each sample corresponded to a unique pair of barcodes. All 32 PCR products were mixed well at a 1: 1 molar ratio and subjected to NGS on an Illumina HiSeq 2500 PE250. The 32 gRNA vectors were also mixed well at a 1: 1 molar ratio and transfected together with BE3 vector. The targeted genomic regions were amplified using site-specific primers with added barcodes and subjected to NGS on an Illumina MiSeq PE300. Control samples were sequenced in the same way, except that they were only transfected with the BE3 vector.

### NGS data analysis

The sequencing reads generated for each sample were first filtered using the Trimmomatic tool with the parameters “LEADING:3 TRAILING:3 SLIDINGWINDOW:4:15 MINLEN:50” to remove low-quality bases at the ends of each read and to truncate reads containing consecutive bases with an average quality score below 15. Using different barcodes, we extracted the sequencing reads for each sample using shell script. We removed unedited reads at gRNA-target sites (20 bp) and collected reads with C:G to T:A conversions at gRNA-target sites (20 bp). The efficiencies of all 32 gRNAs were analyzed by counting the proportion of reads with conversions among all clean reads for each sample. All the above data-processing steps were performed using shell script. We also analyzed the efficiency of each C:G substitution for each gRNA in the same manner. A heatmap was prepared using R software. To determine the indel frequencies, sequence-alignment files in SAM format were generated using sequencing reads and reference sequences via bowtie2. Reads that contained an insertion or a deletion (according to the SAM files) were considered to represent indels.

### Data availability

NGS data have been deposited in the NCBI database under accession code PRJNA434087. Plasmids and cell lines in this study are available upon request. The authors state that all data necessary for confirming the conclusions presented are represented fully in this article.

## Acknowledgement

This work was supported by grants from the National Natural Science Foundation of China (No. 31530071).

